# Temporal sequence geometry enables odor recognition and generalization

**DOI:** 10.64898/2026.01.20.700611

**Authors:** Jonathan V. Gill, Mürsel Karadas, Shy Shoham, Dmitry Rinberg

## Abstract

Neural activity sequences are observed throughout the brain, yet their computational roles remain elusive. In mammalian olfaction, olfactory bulb mitral and tufted cells (MTCs) encode odors with precisely timed activity patterns that tile the respiration cycle. While animals can identify odors independently of concentration within the first 100 milliseconds of inhalation, the structure governing these sequences and the role of activity extending beyond this early window remains unclear. Here, using 2-photon calcium imaging with sub-sniff resolution, we show that odor-evoked MTC sequences propagate as wavefronts through a low-dimensional odor tuning space, where timing is predicted by the tuning similarity between neurons rather than physical location. While early portions of these sequences are concentration-invariant, providing a stable anchor for odor identity, later portions systematically co-activate similarly tuned MTCs across odors, tracing the geometry of the tuning manifold. We propose a role for this later sequential activity in training the piriform cortex to learn perceptually generalizable odor representations. Using a model of Hebbian learning through sequences (HeLSeq), we demonstrate that sequential activity can reinforce synaptic connections from similarly tuned MTCs onto common piriform cortical neurons, enabling rapid generalization to novel odors from the earliest moments of inhalation. These findings support a geometric view of activity sequences and establish a general principle by which temporal sequences scaffold unsupervised manifold learning between brain networks.

## Introduction

Similar to other sensory systems, mammalian olfaction faces a tradeoff between discriminability and generalization^1^. Animals need to identify and discriminate odors independently of their concentration, and at the same time generalize across similar odors^2^. For example, when smelling a new odor like lychee, humans and other animals can quickly assess that it is more similar to other familiar stimuli, such as apricot and grape, than to others, like banana. We investigated this computation in the mouse, whose olfactory pathway is shallow, with only two synapses from the nose to the cortex, and genetically accessible, allowing recordings across stages of the circuit. Odor processing begins when a mouse inhales, as odor molecules are sampled by a set of broadly tuned receptors expressed by olfactory sensory neurons (OSNs) in the nasal epithelium. OSNs expressing the same receptor type converge their axons to form glomeruli with the dendrites of mitral and tufted cells (MTCs) in the main olfactory bulb (OB)^3–5^. Glomerular activity patterns are then reshaped by broad inhibitory recurrence to form the output of the OB through the MTCs before sending this activity to downstream areas, like the piriform cortex^6–10^.

A striking feature of MTC odor responses is their precise temporal patterning across the respiration cycle (sniff). Inhalation of odors activates specific sets of MTCs in discrete time windows with a temporal precision of ∼10 ms, at latencies that tile the sniff cycle (∼330 ms)^11,12^. While rodents can discriminate between odor stimuli using only the earliest portion of this activity (<100 ms)^13–16^, it is unclear how activity in this time window is structured to permit rapid discrimination. Further, the role of activity evolving over the later portion of the sniff is still unknown.

## Results

### Monitoring sub-sniff dynamics across MTCs

Here, to observe odor specific sequences and study their properties we recorded hundreds of simultaneous MTC responses to many odors and concentrations using the fast calcium indicator jGCaMP8f^17^ with sub-sniff temporal precision (**Fig. 1**). Expression was restricted to MTCs by injecting the cre-dependent virus AAV5-syn-FLEX-jGCaMP8f-WPRE into the dorsal OB of Tbx21-cre (Tbet-cre) transgenic mice^18^ and mitral cell layer activity was recorded using 2-photon calcium imaging (**Fig. 1a-c**). We presented a battery of monomolecular odorants (8 odors, 2 concentrations; 5% and 0.5% saturated vapor pressure (SVP)) to awake, head-fixed mice using a custom olfactometer while simultaneously measuring respiration to identify the timing of the first sniff after odor delivery (**Fig. 1d**, **Fig. S1**).

**Figure 1.**
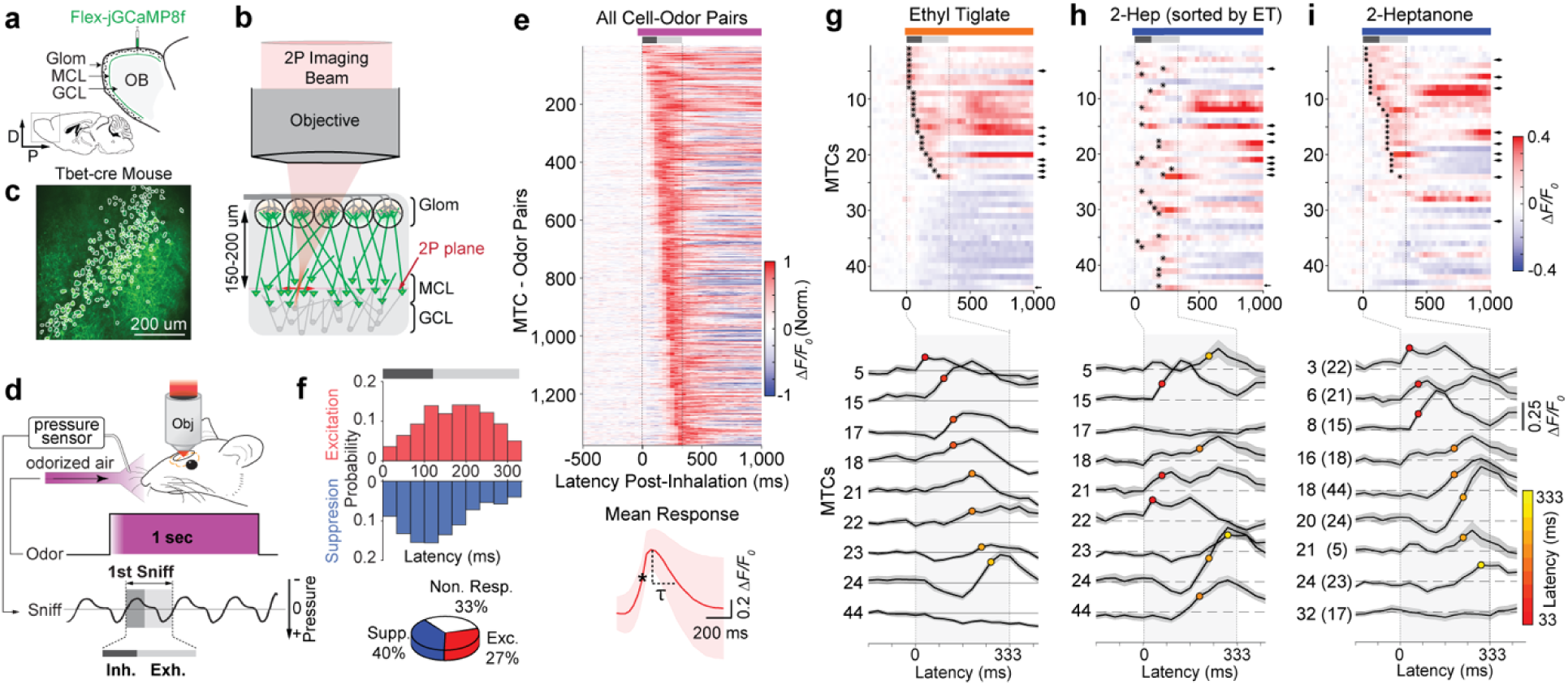
Imaging MTC odor response sequences. **a,** Viral injection and expression of jGCaMP8f targeted to the mitral cell layer (MCL) of the OB in Tbet-cre mice. Glom = Glomerular layer, GCL = granule cell layer. **b,** 2P Imaging of the MCL. Green triangles are MTCs; gray circles are granule cells. **c,** Fluorescence image of jGCaMP8f expression in the MCL (mean of 200 images). Outlines are regions of interest (ROIs) targeting putative MTCs (199 MTCs). **d,** Sniff-triggered odor delivery. The sniff signal was monitored with a pressure sensor placed in front of the nose and odor delivery was initiated during exhalation. The timing of the first sniff was recorded and used to align subsequent analyses. Here and further, the sniff interval is demarcated as a gray bar, with inhalation as a dark gray bar. **e**, *Upper*: Normalized mean fluorescence across excitatory responsive cell-odor pairs aligned to the onset of inhalation (1,376 cell-odor pairs, 8 odors, 5% SVP, 627 MTCs, mean of 10-20 trials normalized to peak response within 333 ms from inhalation). *Lower*: Mean normalized fluorescence across cell-odor pairs aligned to the onset of excitatory response (mean ± 1 standard deviation). Onset time was the first frame exceeding 4 std of the 1 s pre-inhalation baseline (marked *). 𝜏 =160 ms half-decay time. **f**, *Upper*: Histogram of response onset times for excitation (red) and suppression (blue) (suppression onset times were the first frame where mean fluorescence fell below -2 std of the 1 s pre-inhalation baseline, n=5 mice, 8 odors, 5% SVP, 627 MTCs). *Lower:* Percentage of cell-odor pairs exhibiting excitation, suppression, or no response (1,376 excited, 2,005 suppressed, 1,635 non-responsive). **g, h, i,** *Upper:* Mean response across MTCs (n = 44, 15 trials) for one example odor (**g** – ethyl tiglate, **h** & **i** - 2-heptanone) sorted by onset latency for **g** & **i** of the same odor and **h** for onset latency of different odor - ethyl tiglate. *Lower:* Example MTC responses (9 MTCs, mean ± 1 SEM) from the upper panels marked by arrows. Onset latencies are denoted using colored circles.

By aligning odor responses to the onset of inhalation, we observed thousands of cell-odor pairs with response latencies that tiled the sniff cycle (∼333 ms duration, 10 frames at 30 fps, n=1,376 cell odor pairs, 8 odors, 5% SVP, 5 mice) (**Fig. 1e**). All odors evoked selective excitation and suppression in proportions comparable to those found in previous studies using extracellular electrophysiology^12^ (**Fig. 1f**). Suppression was biased toward the late-inhalation period, and excitatory responses were distributed across both inhalation and exhalation (**Fig. 1f**). We observed that excitatory responses activated and decayed in less than one respiration cycle (mean half-decay rate = 160 ms), permitting exploration of temporal dynamics within a single sniff, in contrast to previous imaging studies which only considered activity on the order of several seconds (many sniffs), limited by the previous generation of slower calcium indicators^19^ (**Fig. 1e**, **Fig. S2**).

To explore how odor responses evolved across a population of simultaneously recorded neurons, we ordered mitral cells by activation latency for a single odor (ethyl tiglate, 5% SVP, n = 1 example mouse, mean of 15 trials, **Fig. 1g**). Excitatory responses were distributed across the first sniff, revealing an odor specific sequence. We observed that the response profile reorganized for a different odor (2-heptanone, ordered by ethyl tiglate, 5% SVP, same mouse, mean of 15 trials, **Fig. 1h**), and that by re-ordering the neurons by latency to this second odor, we again revealed an odor specific sequence that unfolded across a different, but overlapping, set of neurons (**Fig. 1i**). These results demonstrate that odor-evoked sequential activity can be resolved within a single sniff, allowing us to explore the structure and role of these sequences across odors and concentrations.

### Geometric representation of odor tuning by MTCs

What determines the ordering and structure of odor specific sequential activity? To explore this organization, we first estimated the odor tuning of individual MTCs by calculating their average response to each odor within the first sniff and combined these to form an ‘odor-tuning vector’ for each MTC (**Fig. 2a**, one example mouse, mean of 333 ms, 15-20 trials per odor and concentration, n = 44 MTCs). We included both excitatory and inhibitory responses of MTCs across odorants as components of the odor tuning vectors. Next, we estimated the pairwise correlations between MTC odor tuning vectors and found a graded, yet modular structure to MTC tuning correlations, with some pairs being highly correlated, others being anti-correlated and many displaying a range of weaker correlations (**Fig. 2b**. *r* = Pearson’s correlation, n = 44 MTCs).

**Figure 2.**
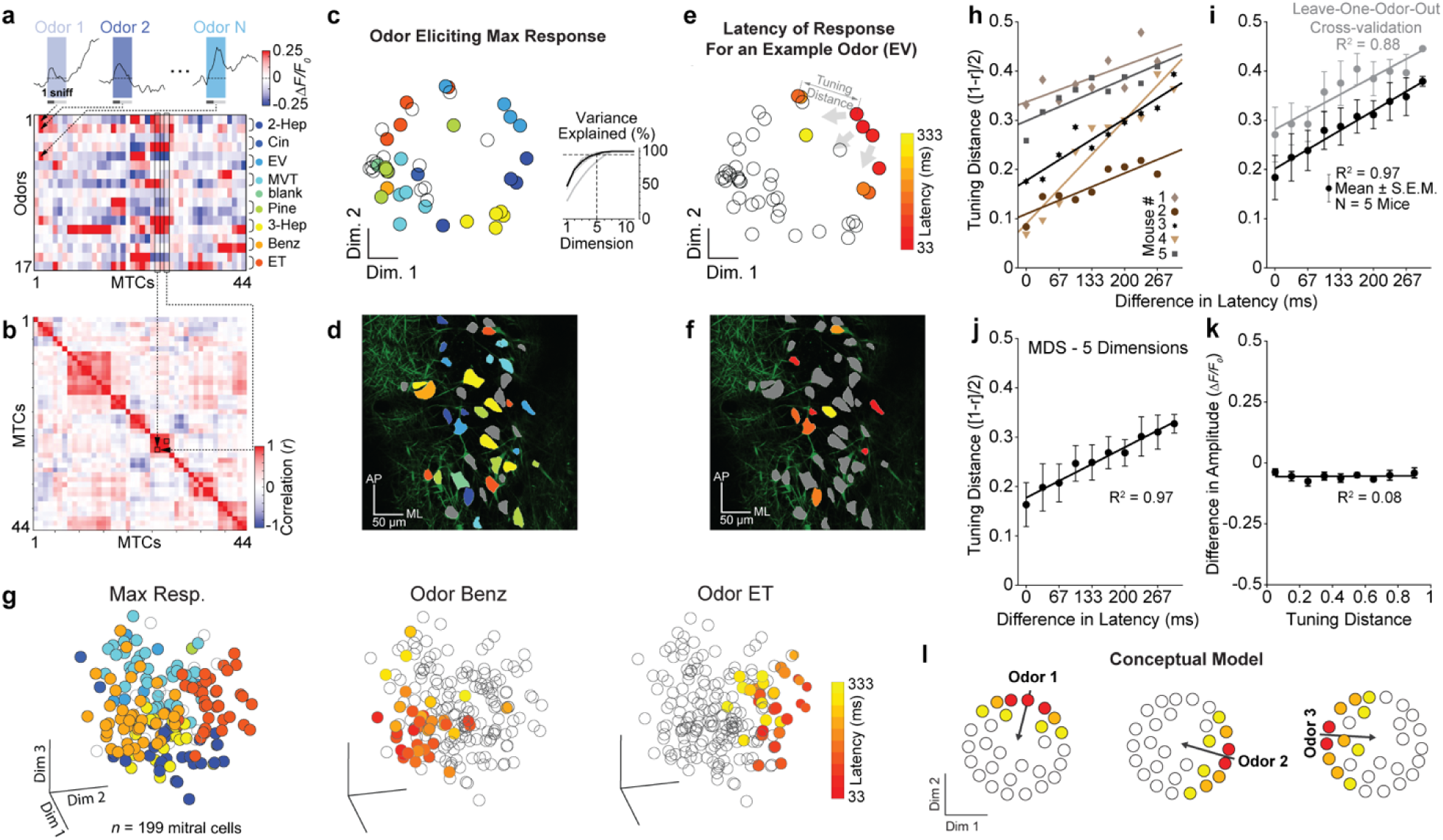
Geometry of odor tuning and sequence propagation. **a.** Odor response matrix for an example MTC field. Odor responses were averaged over a single sniff period (333 ms) and trial repetitions to create a matrix of MTC odor responses (44 example MTCs in 1 mouse, 8 odors at 2 concentrations and blank control, 10-15 trials). For each odor, upper and lower rows were 0.5% and 5% SVP, respectively. *Upper inserts:* examples of individual MTC odor responses. MTCs have been ordered using hierarchical clustering based on pairwise-correlations. **b**. Pairwise correlations between MTC odor tuning vectors (Pearson correlation between the columns of the response matrix in **a**, n=44 MTCs in 1 mouse). One example pairwise correlation is highlighted (MTCs 26 and 28, black boxes). **c.** 2D MDS projection of MTCs from **a** and **b** (circles) positioned by their differences in tuning at high odor concentration (8 odors, 5% SVP and blank control). MTCs are pseudocolored by the odor evoking the maximum response. Empty circles did not have a significant positive response (No resp.). *Inset*: Cumulative percent variance explained as a function of dimensions used for MDS embedding (thin lines = individual mice, black = mean of 5 mice, gray = shuffled control with odor responses randomly permuted for each MTC). **d**. MTC outlines pseudocolored by odor evoking the max response (same MTCs as **c**, but visualized by position in the OB). **e**. Same 2D MDS projection as **c**, but pseudocolored by the latency of odor response to an example odor ethyl valerate (EV) (n=44 mitral cells, mean of 15 trials). **f**. MTC outlines pseudocolored by latency of response to EV. **g**. MDS projection and latency visualization for another mouse’s response to odor benzaldehyde (*Left)* and ethyl tiglate (*Right*) (n = 199 MTCs from 1 mouse, mean of 15 trials, color bar as in **e**). **h**. Tuning distance as a function of difference in latency for 5 mice (mean of 8 odors, 5% SVP). **i**. Mean tuning distance vs. latency across mice (black, slope = 0.019 ms^-1^, R^2^=0.97, p=2.3x10^-7^, 8 odors, 5% SVP, n = 5 mice, mean +/- SEM) and leave-one-out cross-validation control, where tuning distances were estimated with n-1 odors and the difference in latency was measured for the held-out odor (gray, slope = 0.018 ms^-1^, R^2^=0.88, p=7.0x10^-5^, 8 odors, 5% SVP, n = 5 mice, mean +/- SEM). **j.** Same as **i** with MTC odor responses projected onto the top 5 MDS dimensions (slope = 0.017 ms^-1^, R^2^=0.97, p=2.8x10^-7^, 8 odors, 5% SVP, n=5 mice). **k.** Same as **i**, but plotting the mean difference in amplitude vs. the tuning distance between responsive MTCs (slope = 0.004 ms^-1^, R^2^=0.08, p=0.82. 8 odors, 5% SVP, n=5 mice). **l.** A conceptual model where responses to different odors activate sets of MTCs in wavefront-like sequences coming from different angles.

To visualize this correlation structure in a lower dimensional space, we constructed an MTC tuning space using multidimensional scaling (MDS)^20^, a linear method which seeks to preserve the pairwise distances between MTC odor tuning vectors. Here we could project each mitral cell into a lower-dimensional MTC tuning space where individual axes corresponded to the dimensions of greatest variance in odor tuning. Each MTC tuning vector therefore was embedded based on the similarity of its ‘odor receptive field’ to those of all the other simultaneously recorded MTCs. The positions of the full set of MTCs effectively tiled a subspace that could be described as the odor tuning manifold. Quantifying the variance explained by each dimension of the MTC tuning space, we found that 95% of the variance could be captured by 5 dimensions, suggesting an intrinsically lower dimensional structure (**Fig. 2c**, 8 odors, 5% SVP). Projecting MTCs in a 2D MDS space for visualization, we found a smooth subspace of odor tuning, where MTCs sharing a similar maximum odor response were clustered together and MTCs preferring chemically similar odors were positioned near each other (for example, 2-heptanone and 3-heptanone, which differ only by the position of a carbonyl group) (**Fig. 2c**). This was in stark contrast to the ‘salt-and-pepper’ organization of MTC tuning in physical space (characterized by their x-y positions) (**Fig. 2d**).

### Propagation of sequential activity

Next, we analyzed the propagation of sequences in this MDS space and discovered an orderly organization. Sequences originated in groups of co-tuned MTCs and spread to nearby MTCs, expanding into the tuning space with increasing response latency, resembling wavefront propagation (**Fig. 2e**, n = 44 MTCs projected into two dimensions). However, we did not observe obvious wavefront propagation structure in MTC physical space (**Fig. 2f**). We observed these propagation dynamics in MTC tuning space across odors and animals (see **Fig. 2g**, n = 199 MTCs projected into to 3 dimensions for visualization, 8 odors, 5% SVP). While these examples provide a good visualization of wavefront propagation in low D, how general is this effect in the full dimensional space? To quantify this effect, for each MTC that responded at the earliest latency to a given odor, we computed both the tuning distance and the difference in response latency to every other positively responsive neuron. When we averaged across odors (**Fig. 2h**) and mice (**Fig. 2i**) we found a consistent linear relationship between the tuning distance and difference in latency from the earliest responding MTCs to the later responding MTCs, revealing the organization and rate of sequence propagation (**Fig. 2i**, linear regression, slope = 0.019 ms^-1^, R^2^=0.97, p=2.3x10^-7^, 8 odors, 5% SVP, n = 5 mice). This relationship was preserved when the same analysis was performed using only the top 5 MDS dimensions (those representing >95% of the variance), which demonstrates that the propagation dynamics evolve along the lower-dimensional odor tuning space (**Fig. 2j**, linear regression, slope = 0.017 ms^-1^, R^2^=0.97, p=2.8x10^-7^, 8 odors, 5% SVP, n = 5 mice). This relationship also held when we repeated this analysis including only odors at lower concentration (**Fig. S3a-c**, slope = 0.017 ms^-1^, R^2^=0.73, p=1.6x10^-3^, 8 odors, 0.5% SVP, n=5 mice) and with odors at both concentrations (**Fig. S3d-f**, slope = 0.015 ms^-1^, R^2^=0.84, p=2.1x10^-4^, 8 odors, 5% and 0.5% SVP, n=5 mice).

In the previous analysis, the tuning distance between MTCs was parameterized by the responses to all odors in the odor set and the timing estimation was computed per odor. In order to test how robustly we can estimate the geometric organization of sequences, we performed a leave-one-out cross-validation control, building the tuning embedding space using n-1 odors, and testing the relationship between tuning distance and timing for the held-out odor (**Fig. 2i**). We permuted which odor was held out so that we could estimate the distance vs. timing relationship averaged over only held out odors across all of the mice and found that the slope of the curve was consistent with the previous result (**Fig. 2i**, linear regression, slope = 0.018 ms^-1^, R^2^=0.88, p=7.0x10^-5^, 8 odors, 5% SVP, n = 5 mice), demonstrating that the relationship is robust enough to predict the sequence propagation of odors that we did not explicitly use to build the tuning embedding space.

Another potential caveat is that the propagation can be explained by an explicit relationship between the latency and amplitude of MTC odor responses. This relationship has been seen at the level of the glomerular activity^21^ yet has not been examined with high temporal fidelity at the level of the MTCs, which substantially reshape glomerular inputs. To address this, first we examined the relationship between latency and peak amplitude across all excitatory responses and cell-odor pairs (**Fig. S4).** We found that the median amplitude did not decay monotonically across onset latencies and stabilized by ∼200 ms, strongly suggesting there is not a sufficient relationship between amplitude and timing to explain our result. We further tested this relationship by repeating the same analysis from **Fig. 2i**, except substituting the difference in mean amplitude between MTCs instead of the difference in latency. We found no relationship between the tuning distance and difference in amplitude from the earliest responding MTCs to the later responding MTCs (**Fig. 2k**, linear regression, slope = 0.004 ms^-1^, R^2^=0.08, p=0.82), confirming that the propagation of the odor response wavefront is determined by the latency, and not amplitude of the responsive MTCs.

These results can be summarized by a conceptual model in which different odors activate sets of MTCs in sequences that progress along different directions, and in which MTC activation timing is odor-specific and determined by each MTC’s tuning across many odors (**Fig. 2l**). In this framework, the difference in tuning between two MTCs to a set of odors predicts the difference in their response latencies within the sequence evoked by a given odor.

### Relationship between timing and concentration invariance

While the previous analyses examined MTC responses to odors presented at a fixed concentration, animals in the wild typically experience the same odorants across a broad range of concentrations. How are MTC sequences organized across changes in odor concentration? We presented odorants at a range of concentrations (5%, 1% and 0.5% SVP) and observed individual neurons that either maintained a positive response (*consistent*, **Fig. 3a**, *Upper*) or changed their response sign across concentrations (*inconsistent*, **Fig. 3a**, *Lower*). By observing the full sequence of MTC responses to an example odorant, and maintaining their ordering over changes in concentration, we were able to observe a general principle of their concentration dependence: the earliest responding neurons were consistent across concentrations, while the later responding neurons were more inconsistent (**Fig. 3b**). We quantified this effect across odorants comparing responses at high and low concentrations (5% and 0.5% SVP), and though we observed consistent and inconsistent responses at all latencies, we found a significantly higher probability of observing consistent responses if they occurred within the first 100 ms following inhalation, with the greatest difference appearing in the first 33-66 ms (**Fig. 3c,** n=5 mice, 1,376 MTC pairs, 8 odorants, *Upper*: p=5x10^-12^, Kolmogorov-Smirnov test, *Lower*: p=3x10^-15^ Mann-Whitney U test).

**Figure 3.**
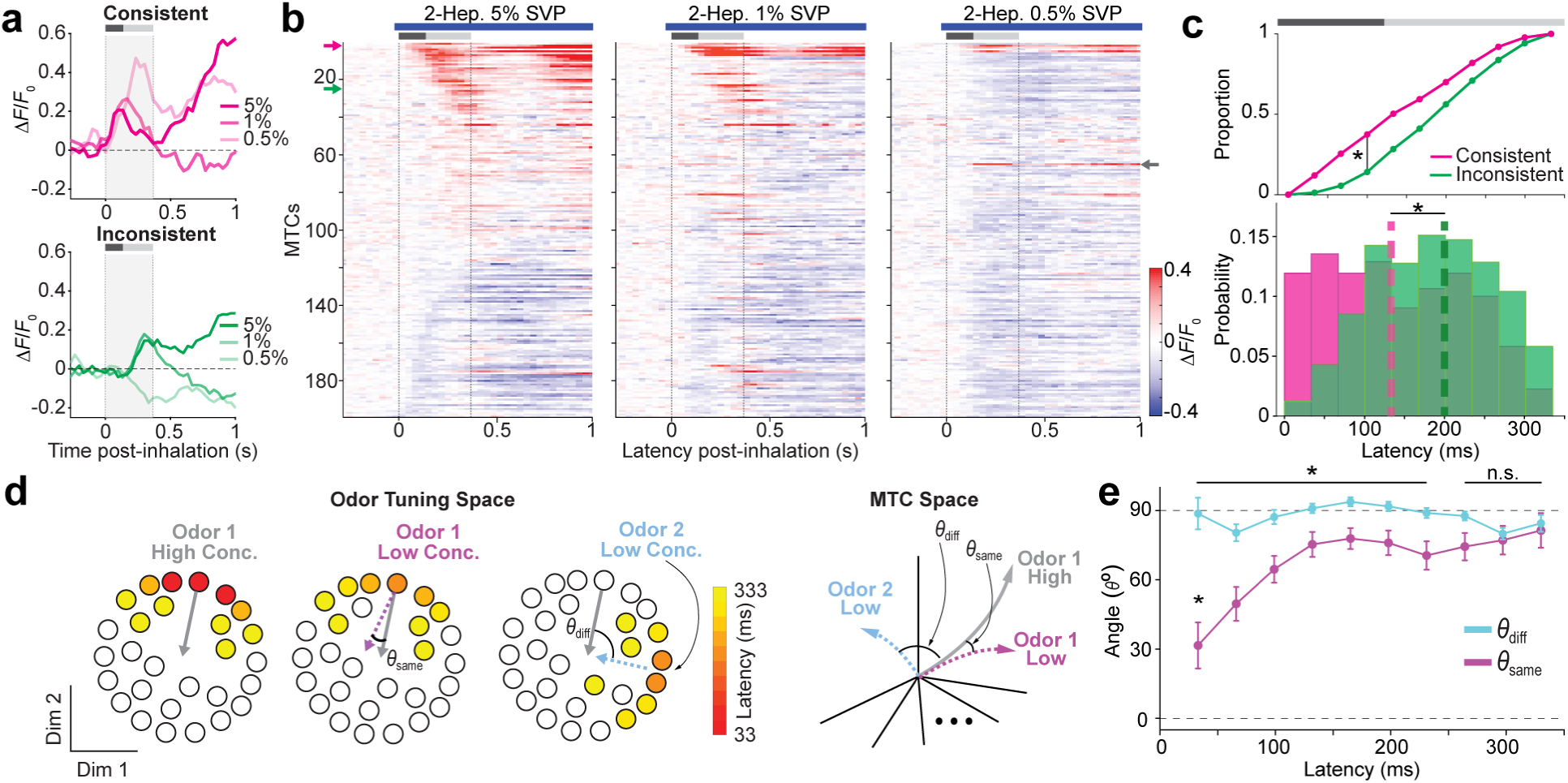
Geometry of odor response sequences across concentrations. **a,** Example concentration consistent (*upper panel*) and inconsistent (*lower panel*) MTC responses to 2-heptanone (2-Hep) at 3 concentrations, 5%, 1% and 0.5% SVP (mean of 10 trials), gray shading and bar indicates 333 ms sniff window). **b,** Mean population response to 2-Hep at high (*left*), medium (*middle*), and low (*right*) concentrations, ordered by onset latency at high concentration (10 trials, n=199 MTCs). Colored arrows denote consistent and inconsistent responsive neurons from **a**. The gray arrow indicates a neuron with a response that emerges at the lowest concentration. **c,** *Lower:* Histogram of the probability of concentration consistent or inconsistent responses as a function of onset latency (1,376 cell-odor pairs, n = 627 MTCs in 5 mice, 8 odors at 5% and 0.5% SVP, p=3x10^-15^ Mann-Whitney U test). *Upper:* Cumulative distributions of data plotted below (vertical black bar = latency of maximum difference, p=5x10^-12^, Kolmogorov-Smirnov test). **d,** *Left*: Conceptual schematic of the angular difference between odor response sequences across concentrations. We measured the angle between the population vectors of the same odor at high and low concentrations (left and middle), or a different odor (right). *Right*: The same analysis can be visualized as comparing the odor response vectors in a space parameterized by responses across MTCs evolving with time from inhalation (latency). **e**, The distribution of angles between the same odor at high and low concentrations (purple) and different odors (blue) as a function of the instantaneous latency from inhalation (1,376 cell-odor pairs, n = 627 MTCs in 5 mice, 8 odors at 5% and 0.5% SVP). The mean of angles between the same odor and different odor population vectors were significantly different from each other from 0-233 ms (p<0.05, two-sample t-tests, corrected for multiple comparisons using false discovery rate (FDR)) and the mean angle for comparisons between concentrations of the same odor were significantly different between the early latency period compared to later latencies (33 ms vs. 133-333ms; p<0.05, ANOVA and Tukey’s test for multiple comparisons).

Importantly, while early responses were more consistent, we found many excitatory responses that appeared only at lower concentrations (see **Fig. 3b**, *Right*: gray arrow highlighting 0.5% SVP response of one MTC). One interpretation is that the population response at low concentrations is not simply a scaled-down version of the population response at high concentrations, but a unique sequence that is concentration dependent. How did changes in concentration affect the geometry of the odor response sequences? A way to quantify this is by constructing population response vectors at high and low concentrations, defined as vectors built from the mean response across all simultaneously recorded MTCs at each latency after inhalation. We measured the angle between population response vectors at high concentration and the population response vectors at low concentration of either the same odorant, or a different odorant (**Fig. 3d**). Two different ways can be used to conceptualize this comparison, either as the angle between the vectors defining the direction of propagation across MTCs in odor tuning space (**Fig. 3d**, *Left*), or comparing the odor response population vectors in a space spanned by the relative activation of all MTCs (**Fig. 3d**, *Right*), which are equivalent with no dimensionality reduction applied. We examined the angle as a function of response latency, comparing the vectors at each time bin across the time course of the odor response (**Fig. 3e**, 0-333 ms, 33.3 ms time bins). We found the smallest angle between response vectors for high and low concentrations of the same odor in the first time bin (0-33.3 ms, mean = 32° ± 10° S.E.M), which evolved to an asymptotic value of ∼76° by 133 ms (33 ms vs. 133-333ms; p<0.05, ANOVA and Tukey’s test for multiple comparisons), while the mean angle between different odors was ∼90° across all time bins (mean = 87.4° ± 3° S.E.M). These results show that response sequences to different concentrations of the same odor begin in a similar set of MTCs and propagate in a similar direction for the first 100 ms, before diverging in the later part of the sniff, ultimately becoming as dissimilar as responses to a different odorant (* = p<0.05, n.s. = p>0.05, all comparisons between groups, two-sided t-test corrected for multiple comparisons using false discovery rate).

### The role of early vs. late sequential activity for guiding perception

Animals can discriminate between odors from the earliest portion of the sniff^13^. What is the potential role of odor sequences beyond the earliest responses? The organization of MTC space in which odors evoke a traveling wavefront-like propagation of activity sets up the condition where if two cells nearby in this space are activated by an odor, they will be co-activated in the same temporal window across different odors (**Fig. 4a**). This nearly coincident activation will occur independently of 1) the direction of the wave propagation and 2) timing in the sniff. This organization potentially creates a robust substrate for learning the correlation between MTCs in areas receiving projections from the OB, like the piriform cortex (**Fig. 4b**). We suggest that this may have a profound effect for establishing perceptual similarities across odors, which we assume is related to the similarity of evoked neural responses (**Fig. 4c**). For example, when smelling a new odor (for example, lychee, **Fig. 4d**), humans and other animals can perceive the stimulus upon the first exposure and assess its similarity to other previously experienced stimuli (e.g., apricot and grape), an ability known as perceptual generalization^2^ (**Fig. 4e)**. Further, given the rapid speed of odorant identification, we posit that the appropriate cortical representation should be formed within <100 ms after odor inhalation^13^, constraining the relevant timing window for generalization to the early inhalation period.

**Figure 4.**
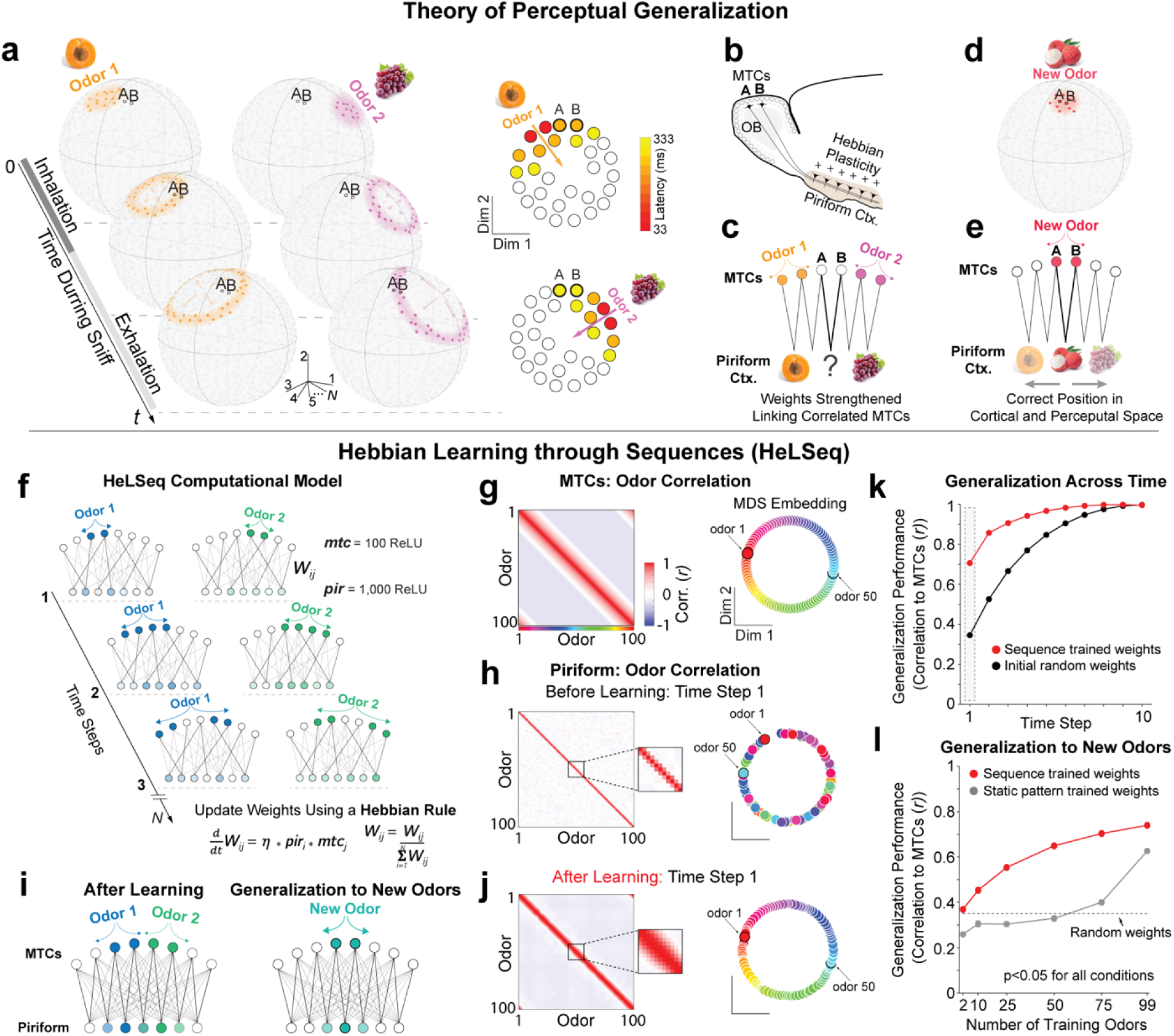
Hebbian learning of readout connections using sequences permits generalization. **a**. *Left:* Illustration of odor response sequences propagating across MTCs positioned on the hull of a hypersphere representing the odor tuning manifold (gray circles). Two MTCs are highlighted (A and B) and co-activated either during the middle (odor 1) or late (odor 2) sniff period. *Right:* A simplified depiction of odor response propagation in 2 dimensions, as in Figs. 2 and 3. **b.** Coactivation of MTCs A and B increases their shared synaptic weights to piriform cortex neurons through Hebbian plasticity. **c.** Exposure to odors 1 and 2 strengthen the shared weights of MTCs A and B before any odor has activated them during early inhalation. **d.** A new odor (lychee) activates MTCs A and B in the early inhalation period. **e.** Shared synaptic weights between MTCs A and B allow the new odor to activate the piriform cortex in a pattern that is correctly positioned between previously experienced odors, allowing for perceptual generalization from the early inhalation period. **f**, Weights between MTCs and piriform neurons are initialized randomly from a half-normal distribution (rectified linear units (ReLU), n=100 MTCs, 1,000 piriform, 0.2 prob. of non-zero weight). Sequences radiate across the MTC layer creating an odor response space of known geometry. Weights between MTCs and piriform neurons are updated at each time step based on a Hebbian rule and normalized to sum to 1 after each odor epoch. **g,** *Left:* The odor-odor correlation for each MTC response pattern averaged over all timesteps (mean of 10 timesteps, 100 odors, Pearson correlation). *Right*: 2D MDS embedding of the odors based on their MTC response correlations. Odor 1 and 50 highlighted. **h,** The odor-odor correlation across piriform neurons before learning at the first timestep (1 timestep, 100 odors, Pearson correlation). *Inset:* correlation structure near the diagonal. *Right*: 2D MDS embedding of the odors based on their piriform response correlations before learning. **i,** *Left:* After learning, weights are enhanced between highly correlated MTCs and individual piriform neurons. *Right:* Presenting an unexperienced odor after training evokes a piriform response pattern that is distinct but overlapping the response to experience odors 1 and 2. **j,** Same as **h**, but after learning. *Inset:* correlation structure near the diagonal. *Right*: 2D MDS embedding of the odors based on their piriform response correlations after learning. **k,** Generalization across odor sequence timesteps for initial random weights and sequence trained weights calculated as the cumulative mean piriform odor response correlation to the mean MTC odor input (mean of 10 network initializations and n = 100 odors). Error bars (not visible) ± s.e.m., p<0.05, paired t-test for all points between trained and untrained performance. Odor-odor correlations in **h** and **j** were included in time bin one performance, highlighted in gray. **l,** Mean generalization performance to new odors, calculated as the correlation between piriform odor-odor correlation structure at the first time step and the mean MTC odor-odor correlation structure. Performance is plotted as a function of the number of odors used for training (40 network initializations per data point, 100 total odors, only untrained odors used for testing). Training was performed either using sequences, or static patterns (all activity in the sequence presented in a single time bin). Dotted line = baseline performance (correlation) with initial untrained random weights. Error bars ± s.e.m., p<0.05, paired t-test for all points between trained and untrained performance.

### A model of perceptual generalization using MTC sequences

To explore how temporally correlated MTC activity could give rise to this perceptual phenomenon, we constructed a biologically plausible network model that makes use of the discovery that similarly tuned MTCs are activated together within a brief time window across multiple odors. In this model, synaptic weights from MTCs to the piriform cortex are initially random, then learned by exploiting the presence of organized sequential MTC activity along with a Hebbian learning rule (here called ‘Hebbian learning through sequences’, or HeLSeq) (**Fig. 4f-l**). We used this model and learning rule to study how correlated MTC activity, even late in odor response sequences, can be used to build a robust representation allowing the piriform cortex to generalize to new stimuli within the first tens of milliseconds after inhaling a novel odor.

For the piriform cortex to ‘know’ about an odor means to have information about the relationship between a specific odor and all other odors. We may assume that this information is initially present in the pairwise odor-odor correlations between MTC odor responses. A requirement for the model is that, after training on some number of odors, we can reproduce the full odor-odor correlation structure of the MTC inputs from the earliest piriform cortical activity. Previous studies have suggested that modeling the synaptic connections from the MTCs to the piriform cortex using sparse, random matrices is sufficient to preserve MTC odor-odor correlations in piriform cortical activity^10,22,23^. However, these models simulated MTC odor responses as static vectors instead of temporally evolving sequences and therefore required the full MTC odor response vector (equivalent of a full sniff) to reproduce odor-odor correlations in the piriform cortex. Therefore, we expected that both random and HeLSeq trained weights would preserve the MTC odor-odor correlations, but that HeLSeq trained weights should reproduce this correlation structure much earlier. This was indeed revealed through the following simulations and analyses.

We constructed a two-layer network of rectified linear units (n = 100 MTCs, 1,000 piriform cortical pyramidal cells) initialized with sparse, random, feed-forward weights (weights were drawn from a positive, half-normal distribution and randomly pruned to achieve 20% non-zero weights (*see Methods*). MTCs were activated by artificial odors in radially sequential patterns that propagated for 10 timesteps (333 ms), which we treated as one sniff. To represent the odor-odor correlations of the MTC inputs, we averaged responses across the sniff for the full set of these patterns (100 odors) and computed their correlation to each other (**Fig. 4g**). By design, the MTC odor-odor correlation space was smoothly varying, with positive correlations representing the overlap between similar odors and negative correlations between dissimilar odors, a shape well described by a ring (**Fig. 4g**, *right panel*). Initially, sparse-random weights led to piriform activation patterns at the first time step that only preserved the strongest correlations (**Fig. 4h**, near-diagonal), and introduced many spurious correlations (**Fig. 4h**, off-diagonal), scrambling odor positions on the ring geometry (**Fig. 4h**, *right panel*). During HeLSeq training, weights were updated proportional to the coactivation of MTCs and piriform neurons and homeostatically renormalized (piriform neuron input weights sum to 1) after each odor epoch (**Fig. 4f**). We found that piriform neurons became selective to highly correlated MTCs (**Fig. 4i***, left panel*), and the population learned to reproduce an accurate, sparsified version of the MTC odor-odor correlations from the first time step (**Fig. 4g,j-k**) that captured the essential features of their geometry (**Fig. 4g,j**; *right panels*). Cumulatively averaging piriform odor responses across time steps revealed that sequence trained weights significantly outperformed initial random weights in reproducing the average MTC odor-odor correlation structure across time steps, especially in the early response period, with random projections taking until the end of the odor epoch to converge to a similar representation (time step 1: Pearson *r* = 0.7 for sequence trained weights, *r* = 0.34 for initial weights; p<0.01, paired t-test for all points between trained and untrained performance).

Because odor response sequences train the synapses between correlated MTCs beyond the earliest responses, we hypothesized that the MTC odor-odor correlation structure could be learned by training the network with less than the full set of odors. To test the generalization ability of the model, we varied the number of odors used to train the network and compared the odor-odor correlation structure of the trained piriform layer to the odor-odor correlations exhibited by the MTCs (**Fig. 4i,l**). Specifically, we compared the odor-odor correlations of the piriform layer at the first timepoint to the odor-odor correlations of the mean MTC responses across the sniff for the held-out odors not used in training. Generalization performance was quantified as the Pearson correlation between the odor-odor correlation structure of these two layers for held out odors. When trained on only a subset of the odors, we found that the trained network could reproduce the space of input correlations better than the untrained network with as few as 2 training odors (**Fig. 4l**, 40 network initializations per data point, error bars are ± s.e.m., p<0.05 for all comparisons). Finally, we tested whether the sequential nature of odor presentations was necessary for perceptual generalization by presenting the same odors as static patterns of MTC activity instead of sequential activation across MTCs. By construction, this manipulation did not affect the odor-odor correlation structure of the MTCs. When trained with the full set of odors presented as static patterns, the MTC odor-odor correlation structure was well preserved across the piriform responses (**Fig. 4l**, 40 network initializations per data point, error bars are ± s.e.m., p<0.05 for all comparisons). However, when the model was trained with fewer odors, the performance fell below the performance of the untrained, random network, demonstrating a lack of generalization (**Fig. 4l**, 40 network initializations per data point, error bars are ± s.e.m., p<0.05 for all comparisons). Taken together, these results demonstrate how HeLSeq permits efficient transmission of representational geometry across layers of processing, beyond what is achievable by only considering neural activity as static patterns.

## Discussion

Precise, reliable sequences of neural activity have been described across diverse brain regions, from sensory and motor areas to those involved in navigation and planning^24–32^. However, despite their ubiquity, the role of sequential activity in neural computation is not always clear. For example, a clear relationship exists when moving stimuli evoke sequences of activation across the retina. In this case the timing and dimensionality is defined by the temporal structure of the stimuli; precise neuronal spiking follows the movement of the stimuli and patterns evolve over two dimensions^33,34^. However, sequential activity is found in many areas encoding sensory, action or decision variables that can only be described with higher dimensions^26,29^. Along what dimensions are sequences organized in this more general context, and what roles do they serve?

The olfactory system, which is thought to encode the high-dimensional chemical space^35–37^, provides a platform to assess the functional role of sequences^38–41^. Here, recent breakthroughs in the development of fast calcium indicators^17^ allowed us to record hundreds of cells simultaneously with sub-sniff resolution, permitting observation of these sequences across many neurons and found wavefront-like propagation in a low-D odor-tuning space spanned by the mitral cells. Our study contrasts with previous attempts, which have been hampered by several constraints. The first is the typically low numbers of simultaneously recorded neurons collected with extracellular electrophysiology. Although this method offers high temporal resolution, recordings in the olfactory bulb usually yield few neurons and may be biased toward cells with high firing rates. The second constraint is that slow calcium indicators traditionally employed are unable to reveal sub-sniff dynamics (**Fig. S2**). The third constraint comes from using a low number of stimuli, or only a single concentration, making it difficult to estimate the differences in tuning between MTCs (but see^35^).

By measuring responses to many odors in many mitral cells we were able to define an odor tuning space inhabited by these neurons. The space is low dimensional, and in this space, cells that respond to odors similarly live close to each other, in contrast to their disordered (‘salt and pepper’) organization in physical space^35^. It is likely that the odor tuning properties of individual MTCs are formed through both a combination of the receptor affinity of their parent glomerulus, as well as lateral and centrifugal inputs which can drastically reshape their responses^21,42–48^. However, it is unknown how lateral, centrifugal and neuromodulatory mechanisms interact to shape the odor tuning space of MTCs and the timing of odor specific sequences observed in this study. We believe that our findings and the approach employed in this study provide a quantitatively rigorous platform with which to address the impact of different circuit mechanisms on bulbar computation.

The precise timing of MTCs with respect to respiration has been observed in previous studies^11,12^. However, the relationship of timing to the tuning properties of the neurons has not been explored. Here we demonstrated that sequences are orderly and predictable from the perspective of activity waves propagating rapidly across MTCs in odor tuning space. Sequential activity originates in a small number of highly correlated MTCs before spreading to neurons more distantly positioned in this space, with an approximately linear relationship between odor tuning similarity and timing. This means that cells nearby in the odor tuning space will also be highly correlated in time, regardless of where they are positioned in this space.

What role does sequential activity play for guiding perception and behavior? Animals use their olfactory system for a range of behaviors, including navigation. Odor concentration can change several orders of magnitude from when a mouse picks up on a faint whiff of odor to when it navigates to the source. Like walking towards a dim light in the dark, the spot grows on your retina, activating a larger patch of photoreceptors, indicating you’ve moved closer. Similarly, inhaling an odor of higher concentration activates more OSNs and subsequently a larger patch of MTCs, except instead of being positioned in retinotopic space, they are positioned in odor-tuning space. This is where the analogy ends, as defining the ‘center’ of the patch in high dimensional odor-tuning space isn’t as simple as it is for vision, and keeping track of the odor identity across concentrations has the potential to prove challenging for the mouse. Here we find that a unifying feature of odor response sequences across concentrations is that the earliest part of the response tends to remain consistent, while the later part of the response can vary substantially. While the early portion of the code is not necessarily *identical* across concentrations, it is highly similar, serving as an anchor for odor identity across changing conditions. This feature likely underlies the ability of animals (including humans^49^) to make judgments of odor identity very rapidly (within <100 ms)^13–16,50^, while considering relative intensity as a mostly separate perceptual dimension.

Building on behavioral results demonstrating rapid identification of odors, as well as the observation that odor specific sequences extend well beyond this initial <100 ms window, we propose a novel role for this later activity in establishing MTC wiring to the piriform cortex that could serve both rapid odor discrimination and perceptual generalization. We posit that through the course of passive odor exposure, piriform cortical neurons may learn to become ‘coincidence detectors’, becoming sensitive to highly correlated MTCs. This is established by modifying initially random MTC-to-piriform synapses using Hebbian plasticity^51^, which these synapses display during development and to a lesser degree adulthood^52,53^ (though see^54^). The end result is that odors evoking similar responses in MTCs are mapped to overlapping patterns in the piriform cortex *from the first tens of milliseconds*.

In an effort to explain how MTC sequences can guide the formation of generalized representations in the piriform cortex, we propose a simple learning rule, HeLSeq, that generalizes Hebbian learning to sequences of activity. The HeLSeq algorithm is biologically plausible, numerically stable, and robust to overfitting, as demonstrated by our model’s high generalization accuracy with few training examples. We posit that HeLSeq is one example within a range of solutions whereby networks of neurons can learn the shape of the relevant activity manifold of their inputs from upstream regions after experiencing only a few stimuli. The key concept is that, like the swing of a tennis racquet, neural activity extends beyond the initial, behaviorally relevant period (striking the ball), evolving along a trajectory that traces the shape of the manifold (the follow-through), and permits learning the mapping between potential patterns of activity and their relationships to each other (e.g., how racquet angle relates to where the ball lands)^55^. This way, unexplored portions of the potential space of neural activity can be rapidly mapped to their meaning, permitting flexible behavior in novel situations (e.g., encountering a new odor, returning a difficult serve, etc.).

These findings have a striking resemblance to phenomena found in other circuits and organisms. For example, previous literature on retinal waves proposed a similar mechanism by which spontaneous activity waves travel across retinal ganglion cells in developing animals, correlating the activity of nearby neurons, and reinforcing their shared synaptic projections to the LGN^56,57^. This process is ultimately responsible for refining retinotopic maps and ocular dominance columns in V1, establishing the functional architecture of the visual system prior to gaining visual experience^58^. Similarly, the precise temporal sequences of neural activity observed during singing in juvenile zebra finches have been implicated in shaping the synaptic connections between areas HVC and RA, coordinating song generation circuits before they crystalize in adulthood^59^. Temporal activity in the mouse OB may play a similar role to these previously discovered mechanisms with a key difference: the OB produces new sequences throughout the life of the animal whenever the mouse experiences new odorants. This raises the intriguing possibility that OB temporal activity may shape and reshape cortical representations through development and into adulthood. This would allow animals to adapt to different chemical environments, like those imposed by changing seasons, migration, and parenthood, and potentially explain paradoxical observations like the presence of both tuning stability and representational drift in the PCx^60^. Observing odor evoked activity in both the OB and PCx over time may resolve these questions in future studies.

Our results have broad implications for defining the role of sequential activity in how stimuli are encoded and communicated between sensory areas. We propose and provide evidence that the early and late parts of these odor specific activity sequences may serve different roles: the early part permitting proper identification of odorants and the later part further setting up the odor representation in the cortex that reflects the perceptual relationships between similar odors. Further, the technical and conceptual contribution of the approach applied here is likely to be broadly applicable to the study of other systems in which information is encoded at fine spatial-temporal scales.

## Acknowledgments

The authors thank the members of the Rinberg and Shoham labs for their assistance and multiple discussions. Special thanks to S. Toole for technical assistance and P. Veeramreddy for managing the mouse colony. This work was supported by the NIDCD grants (R01DC022320 to D.R. and S.S.) and NIH BRAIN Initiative Grants (U19NS107464 to D.R. and S.S., and U19NS112953 to D.R.), J.V.G. was supported by the Leon Levy Postdoctoral Scholarship in Neuroscience.

## Author contributions

Conceptualization: J.V.G., S.S., D.R.; Data curation: J.V.G., M.K.; Formal Analysis: J.V.G.; Funding acquisition: J.V.G., S.S., D.R.; Investigation: J.V.G., M.K., S.S., D.R.; Methodology: J.V.G., M.K., S.S., D.R.; Project administration: D.R.; Software: J.V.G, M.K.; Supervision: S.S., D.R.; Writing: J.V.G., M.K., S.S., D.R.

## RESOURCE AVAILABILITY

### Lead Contact

Further information and requests for resources should be directed to and will be fulfilled by the Lead Contact, Dmitry Rinberg.

### Materials Availability

This study did not generate new unique reagents.

### Data and Code Availability

The datasets generated and/or analyzed during the current study are available from the corresponding authors on reasonable request.

## METHODS

### Mouse lines

Male and female mice between 2 and 4 months old were used in all experiments and handled in accordance with institutional guidelines. All experiments were performed using hemizygous Tbet-cre^18^ (Tbx21-cre, Stock No: 024507, Jackson Laboratories) mice, which were produced through crosses with wild-type (C57BL/6J, 1 mouse, Stock No: 000664 Jackson Laboratories), OMP-ChR2-EYFP^61^ (2 mice), or M72S50-ChR2-EYFP (Olfr545 tm3(Olfr160)Mom)^62^ (2 mice). No differences between mouse lines were observed while performing or analyzing recordings from the mitral cell layer. Animals were kept in a climate-controlled vivarium operating on a reverse 12-hour light schedule (lights on at 20:00 h). Mice were group housed until surgical implantation, after which the animals were housed singly. All procedures were approved under NYU Grossman School of Medicine institutional animal care and use committee (IACUC) protocol 161211-02.

### Surgical preparation

Mice were anesthetized with isoflurane during viral injections and surgical implantation (2.0% during induction, 1.5% during surgery). A circular craniotomy was performed to expose both hemispheres of the dorsal olfactory bulb (3 mm craniotomy extending from the rostral rhinal vein to the naso-frontal suture, centered on the midline) using an air-driven dental drill (Midwest Tradition, FG 1/8 drill bit). The adeno-associated viral (AAV) vector encoding the calcium indicator jGCaMP8f (AAV5-Syn-Flex-jGCaMP8f-WPRE-SV40, Addgene #162379-AAV5) was injected bilaterally using a stereotactic syringe pump (World Precision Instruments Inc.) at a rate of 0.1 µl min^-1^ (600 nl per hemisphere, 400-600 µm deep). Following injection, a cranial window was implanted replacing a circular piece of skull by a glass coverslip (3 mm diameter, Warner Instruments) that was secured in place using a mix of self-curing resin (Orthojet, Lang Dental) and cyanoacrylate glue (Krazy Glue). A custom 3D-printed headpost^63^ was placed around the cranial window and affixed to the skull using C&B Metabond dental cement (Parkell). Each animal recovered for at least 10 days prior to experiments and 2P-imaging was performed 3-4 weeks after virus injection.

### Odor delivery

Odors were prepared and delivered using a custom three-cassette air-dilution olfactometer, based on Nakayama & Rinberg (2022) with slight modifications (**Fig. S1**). A mass flow controller (MFC; Alicat MC-1SLPM-D/5M/5IN) maintained a total clean airflow of 1 L/min. Each cassette contained an odor-line MFC (0–100 mL/min; Alicat MC-100SCCMD/5M/5IN), two inline Teflon four-valve manifolds (NResearch, 225T082), a three-port bypass valve (NResearch, TI1403270), and eight odor vials. Odors were stored in amber volatile organic analysis vials (Restek, 21797). When idle, air from both the carrier and odor-line MFCs flowed through the bypass valve, totaling 1,000 mL/min, and was routed through a final valve (two 3-way valves; NResearch, SH360T042) to an exhaust line. Simultaneously, clean air was delivered through a separate line at ∼1,000 mL/min to the odor port, with flow reduced to ∼500 mL/min via partial vacuum. To present an odor, the bypass valve of a selected cassette was closed while two odor valves opened, directing air through the vial headspace and mixing it with the clean airflow—thus producing an air-diluted odor. Odor concentration was controlled by adjusting the odor-line MFC. For example, setting it to 100 mL/min produced a high concentration. After allowing at least 2 seconds for flow stabilization, the final valve switched to direct the odorized air to the odor port with <40 ms latency. At stimulus end, the valve returned to clean air delivery.

Because liquid-phase dilutions do not reliably predict gas-phase concentrations^64^, all odor concentrations were adjusted in the air-phase using a serial diluter composed of two MFCs. For instance, to achieve a 10× dilution, both vacuum and air MFCs were set to 900 mL/min. (https://github.com/olfa-lab/olfactometry/tree/master/device_firmware/serial_repeater_teensy).

All odors were obtained from Sigma-Aldrich and 0.5-1 mL of each odor was placed in a 45 mL vial. The following odors were used in all experiments:

**Table 1.**

| Odor name | Abbreviation | CAS Number | Air dilution range |
| --- | --- | --- | --- |
| Benzaldehyde | BENZ | 100-52-7 | 0.05 – 0.005 |
| Methyl Valerate | MTV | 624-24-8 | 0.05 – 0.005 |
| Ethyl Valerate | EVT | 539-82-2 | 0.05 – 0.005 |
| 3- Heptanone | 3-HEPN | 106-35-4 | 0.05 – 0.005 |
| Ethyl Tiglate | ETG | 5837-78-5 | 0.05 – 0.005 |
| (+)-a-Pinene | PINE | 7785-70-8 | 0.05 – 0.005 |
| 2-Heptanone | 2-HEPN | 110-43-0 | 0.05 – 0.005 |
| Trans-Cinnamaldehyde | CINALD | 14371-10-9 | 0.05 – 0.005 |

### Closed-loop odor presentation

Timing of odor delivery was controlled in a closed-loop manner relative to respiration phase. Respiration was monitored using a pressure transducer coupled to a custom Teflon ‘odor port’ which continuously passed clean air or odorized air over the mouse’s nostrils at a rate of ∼0.5 L/min^13^. Pressure change relative to atmospheric pressure was measured using an externally mounted pressure sensor (24PCEFJ6G, Honeywell) positioned in front of the animal’s nares. This pressure signal was amplified (AD620, Analog Devices), and a Schmitt (dual-threshold) trigger was used to define inhalation and exhalation onsets in real-time on an Arduino microcontroller. Once a trial was initiated, after flow was stabilized (2 s), the final valve was switched by exhalation onset and odor was delivered to the odor port. The system then detected the first inhalation onset which was used to align analysis of the simultaneously collected imaging frames (**Fig. S1**).

### Imaging

Imaging was performed on two custom multiphoton microscope systems^21,65,66^ based on the MIMMS 1.0 and MIMMS 2.0 designs (HHMI Janelia Research Campus, Ashburn, VA). Two-photon fluorescence of jGCaMP8f was excited at 930 nm using a mode locked, 80 MHz repetition rate, femtosecond-pulsed, Ti:Sapphire laser with dispersion compensation (InSightX3, Spectra-Physics, Mountain View, CA). The beam was relayed and magnified by a telescope (scan lenses, *f* = 35 mm or *f* = 50 mm, and tube lens, *f* = 200 mm) to the back-aperture of a 16×/0.8-NA water immersion objective lens (Nikon). Images were acquired at 30 Hz using resonant-galvanometer raster scanning (Cambridge Research). Emitted photons were reflected by a dichroic mirror (DM1, FF705-Di01, Semrock, or ZT575/140dcrb-uf1 custom laser bandpass reflective dichroic, Chroma) and separated to either green or red channels via a dichroic mirror (DM2, 565dcxr, Chroma) and fluorescence was detected using GaAsP photomultiplier tubes (H10770PB-40, Hamamatsu). Images were digitized and recorded using ScanImage software^67^ (Vidrio Technologies). Imaging power was adjusted per FOV in the range 35-70 mW. The stability of the imaging and head-fixation system permitted imaging of the same FOV over multiple days in longitudinal experiments (see Online Motion Correction).

### Online motion correction

For all imaging experiments we implemented an online motion correction method in order to target the same neurons within and across sessions. In these experiments, the FOV was first aligned to a reference image manually, then the position was fine-tuned automatically using a custom designed closed-loop algorithm^65^ implemented as a module within ScanImage software^67^ (Vidrio Technologies). This algorithm attempted to minimize the difference between the reference image and the FOV by iteratively moving the microscope stage (Sutter 285 or Thorlabs PLS-XY) to reduce the residual displacement computed using a rigid motion correction package (NoRMCorre, Flatiron Institute^68^). The optimization typically converged within 10-15 s once the residual displacement vector was reduced to <0.5 µm in magnitude. In addition to aligning the FOV across days, we performed this routine between consecutive blocks (60 trials, 6-9 minutes) during each imaging session to minimize the effect of slow *x*-*y* drift due to brain and microscope motion, therefore ensuring the imaged neurons remained consistent throughout each session. We monitored for drift in the *z* dimension as well, which was manually corrected using the reference image between blocks (60 trials, 6-9 minutes) if necessary, though displacement was typically small (∼3 µm in a 1.5 hour session).

### Image processing and analysis

Data analysis was performed using custom-written software in ImageJ (NIH) and MATLAB (MathWorks). Images were aligned to a session-averaged template image using a non-rigid motion correction package^68^. Cellular regions of interest (ROIs) were manually drawn using the jGCaMP8f (green) signal, and mean fluorescence time-courses were extracted. ROI selection utilized both mean and maximum intensity projections from 3-5 blocks of trials per session (5-10 min/block). As neurons could be more elaborated in higher or lower z planes, to ensure segmentation of cell somas we also used a *z*-stack of at least 100 µm in depth centered on the focal plane for ROI selection.

### Activation onset latency

MTC jGCaMP8f fluorescence was sampled at approximately 30 Hz (corresponding to 33.33 ms bins). To determine activation onset latency, we computed *ΔF/F_0_*, with *F_0_* defined as mean baseline fluorescence signal *F_0_* acquired during the 1 second period preceding odor inhalation. Neuronal responses were calculated as *ΔF/F_0_* fluorescence signals for each trial (10-20 trials per odor and concentration), and we averaged *ΔF/F_0_* across trials in a 10-frame window (333.33 ms) aligned to the first frame occurring after inhalation onset for each neuron and odor condition (**Fig. S1**). A neuron was considered responsive if it showed a significant difference in its average fluorescence signal in a 333.33 ms window immediately after inhalation compared with 1 s baseline fluorescence immediately preceding inhalation (*p* < 0.05, two-sample Student’s t-test). Significant responses were then classified as either excitation or suppression based on the sign of the sniff-averaged response. Onset latency for excitatory responses was defined as the first time point at which the *ΔF/F_0_* exceeded 4 standard deviations above its baseline, and -2 standard deviations for suppressive responses. The standard deviation was computed independently for each neuron based on its pre-stimulus baseline fluorescence fluctuations, and responsiveness criteria was applied uniformly across all MTCs. Neuronal responses that did not meet these criteria were excluded from latency analyses as response latency could not be accurately determined.

### Odor tuning distance and dimensionality reduction

To determine odor stimulus tuning, we averaged *ΔF/F_0_* across trials in a 10 frame window (333.33 ms) aligned to inhalation onset for each neuron and odor condition and constructed a matrix of these sniff-averaged responses in the form [odors x neurons]. Each column corresponded to an ‘odor tuning vector’ comprised of the sniff-averaged odor responses for each MTC. To calculate odor tuning distances between MTCs, we first calculated the Pearson correlation coefficient (𝑟) between all sets of tuning vectors, creating a correlation matrix of the shape [neurons x neurons]. We then converted this correlation matrix into a distance matrix, bounded between 0 (no difference in tuning) and 1 (opposite tuning) by computing 𝐷 = (1 − 𝑟)/2. For visualization of pairwise MTC odor-tuning distances, we embedded 𝐷 in a reduced dimensional space using classical (metric) multi-dimensional scaling (MDS)^20^. Analyses were performed either on raw distances D (**Fig. 2h,i,k**, **Fig. S3a,b,d,e**), or distances following MDS embedding (**Fig. 2j**, **Fig. S3c,f**). For distances in MDS embedding space, we first computed Euclidean distances between points, then re-scaled the Euclidean distances to bounded distances (1 − 𝑟)/2 for comparison between conditions.

### Comparison of geometry across odor concentrations

We compared the geometric structure of MTC response sequences evoked by different odors and across concentrations of the same odor. First, we constructed an ’odor tuning space’ from sniff-averaged MTC responses (’odor tuning vectors’) to all odors presented at high concentration (5% SVP), following the same procedure described for odor tuning distance analysis. We then identified the sets of MTCs exhibiting excitatory response onsets at each latency bin (bin size = 33.33 ms, 10 bins total) for both high and low concentrations (5% vs 0.5% SVP) of each odor. For each latency bin, we computed the mean of the odor tuning vectors for the co-active MTCs, defining a ’centroid’ tuning vector that represents the average position of the active population in tuning space at that time point. To assess concentration invariance, we measured the angle between centroid tuning vectors across concentrations of the same odor (high vs low) at each latency. As a control, we also measured the angle between centroid tuning vectors across different odors (high concentration of one odor vs low concentration of each other odor) at each latency. Smaller angles between concentrations of the same odor relative to different odors indicate concentration-invariant structure in the sequence. Angles were averaged across all pairwise comparisons and across mice to produce **Fig. 3e**.

### Computational Modeling

Our network model has two layers of feedforward connectivity that we refer to as the MTC layer 𝒙, and the piriform cortical layer 𝒚, respectively, as shown in **Figure 4**. A fraction of the MTC layer was activated by each odor and odor-odor correlations were calculated to quantify the similarity between all odors. Odors were square pulses applied as inputs to a fraction of MTCs (0.2), starting in a set of two neurons and radiating out in time, such that each time step included the activations from the previous timestep and two additional activations of the neighboring MTCs (**Fig. 4f**). Each odor began from a distinct set of two MTCs, such that odor response sequences could be considered as propagating along a ring from different angles, modeling our experimental findings (**Fig. 4g**). Odor-odor Pearson correlations (*r*) were calculated to quantify the similarity between all odors by taking the average MTC response over 10 time steps. The response of each piriform cortex neuron was a threshold-linear function of the sum of its inputs, 𝑦_𝑖_ = Θ[∑_𝑖_ 𝑊_ij_𝑥_j_ − 𝜃], where Θ(𝑥) = 𝑥, if 𝑥 > 0, and Θ(𝑥) = 0 otherwise. The threshold 𝜃𝜃 = 0.0525 was chosen such that an average of 8-10% of piriform neurons were activated by each odor, matching data from previous studies^22,69,70^.

Before training, 𝑊*_ij_* was initialized with random weights drawn from a half- normal distribution 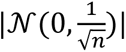, where 𝑛𝑛 is the number of piriform units. We quantified the odor-odor correlations present in the piriform layer across 10 time steps, visualizing the first time step which models the overlap during the early inhalation period (**Fig. 4h,j**). After measuring initial odor-odor correlations, we presented all 100 odors in a randomly permuted order for 1,150 training epochs and updated the weights 𝑊*_ij_* according to a Hebbian learning rule:

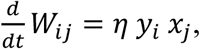

where 𝜂 = 0.0001. After each odor epoch, weights were homeostatically normalized such that ∑*_j_* 𝑊*_ij_* = 1, ensuring competition between weights into each piriform neuron while keeping total input from growing. Generalization across time was computed by first computing odor-odor correlations for piriform responses shaped through random initial weights, or weights trained using HeLSeq at each timepoint of the odor response sequence. We cumulatively averaged piriform responses before computing odor-odor correlations at each time point. We then compared the piriform odor-odor correlations to the sniff-averaged MTC odor-odor correlations using Pearson correlation at each time point. Given the properties of random matrices, we expected that both random and trained weights would preserve the MTC odor-odor correlations, but sequence trained weights should reproduce this correlation structure much earlier, which was revealed through this analysis.

Quantifying generalization performance to new odors was performed by dividing odors randomly into training and test sets, where the number of tested odors was the total odor set minus the training odor set. Training odors were presented in a random order for 1,000 repetitions. After training, odor-odor correlations were calculated from piriform activation within the first time step to the full set of odors, and the odor-odor correlations between the tested odors and the rest of the odor responses was compared to the odor-odor correlations for the same odors from the MTC responses. Similarity between these two representations was quantified using Pearson *r.* To assess the impact of sequential activations for generalization performance, we also trained weights using static patterns which were constructed by summing all activity at the MTC layer for each odor across time steps, and presenting this input in a single time step. The piriform activation threshold was adjusted to 𝜃 = 0.00525 to account for the 10x greater input experienced during static pattern training. After training, odor-odor correlations were calculated from piriform activation within the first time step identically to the analysis for initial random and sequence trained weights.

## SUPPLEMENTARY FIGURES

**Figure S1.**
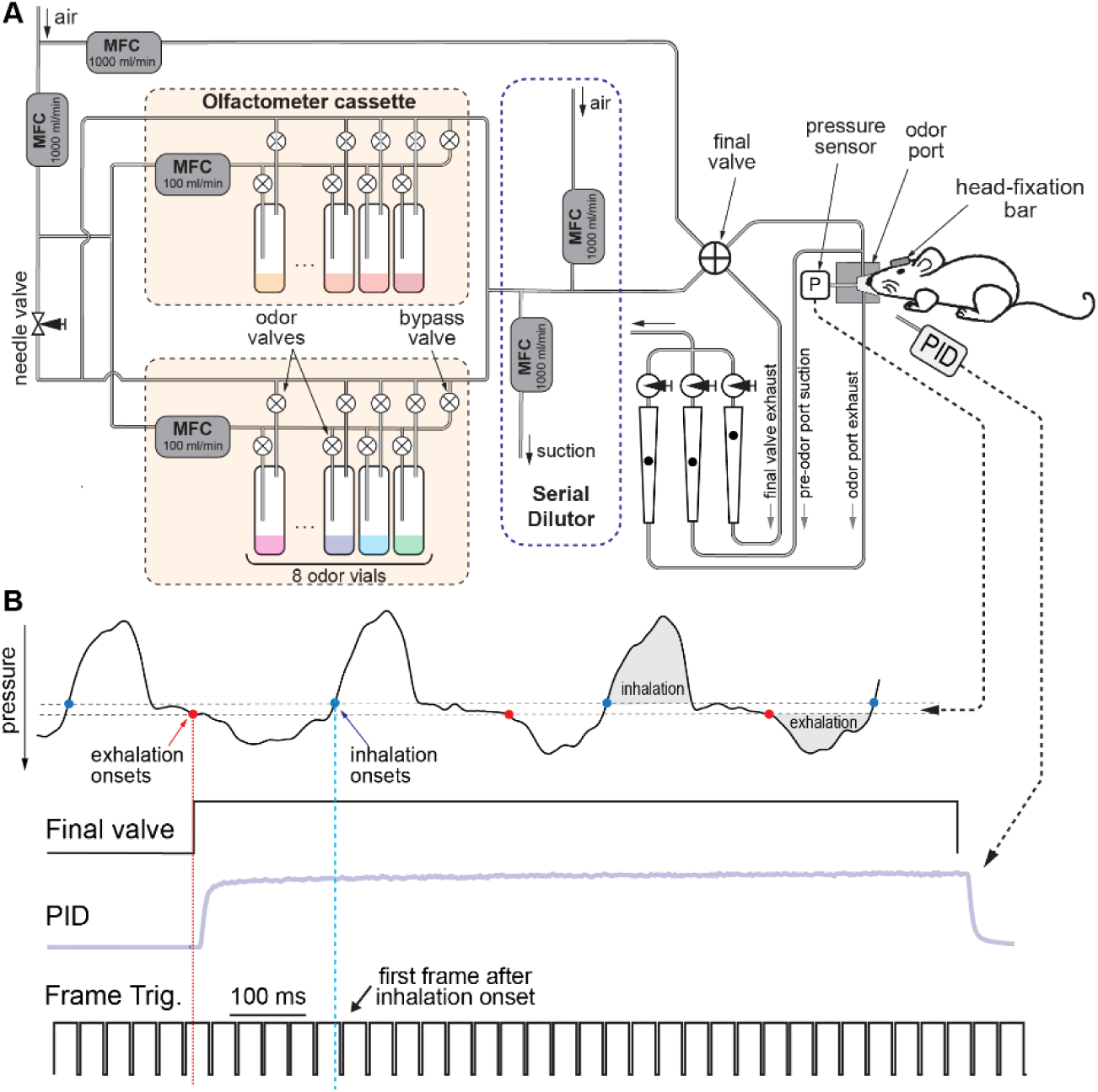
Odor delivery system and alignment to imaging. **A.** Odor stimuli were delivered using a multi-cassette air-dilution olfactometer outfitted with a serial diluter. Each cassette contained eight odor vials, sixteen valves for routing, a 100 mL/min mass flow controller (MFC), and a bypass valve that was default-open. A separate 1,000 mL/min MFC regulated the total airflow through both cassettes. To begin a trial, the two valves corresponding to the selected vial were actuated and the cassette’s bypass valve was closed. Initially, the odorized stream (1,000 mL/min) was directed to the exhaust via the final valve (FV), while a parallel clean-air stream of equal flow rate, metered by an additional MFC, was routed to the odor port. After approximately 2 seconds to reach steady flow, the FV switched so that the odor stream was sent to the port and the clean air was diverted to the exhaust. When the stimulus period ended, the FV returned to its original position, restoring clean air to the port. Odor concentration was set by the relative flow rates of the MFCs, and the integrated serial diluter enabled further attenuation, up to 20-fold, by calibrated withdrawal of odorized air with simultaneous replacement by an equal amount of clean air controlled by two MFCs. Air leaving the FV was sent to both the odor port and a pre-odor port exhaust line in a 50/50 ratio, reducing the total airflow experience by the animal. Air entering the odor port was removed via a matched exhaust line to prevent pressure fluctuations and contamination of the mouse’s surroundings. The port also housed a pressure sensor used both to monitor the animal’s sniffing and to calibrate airflow. **B.** Temporal structure of control signals. Sniffing was continuously monitored via the port pressure sensor. The FV was triggered at the onset of exhalation so that odor concentration stabilized before the first inhalation, verified by measuring the temporal concentration profile at the odor port with a photo-ionization detector (PID). Frame acquisition times from the 2-photon microscope were recorded as TTL pulses (Frame Trig) and used to align data to the first frame after inhalation onset.

**Figure S2.**
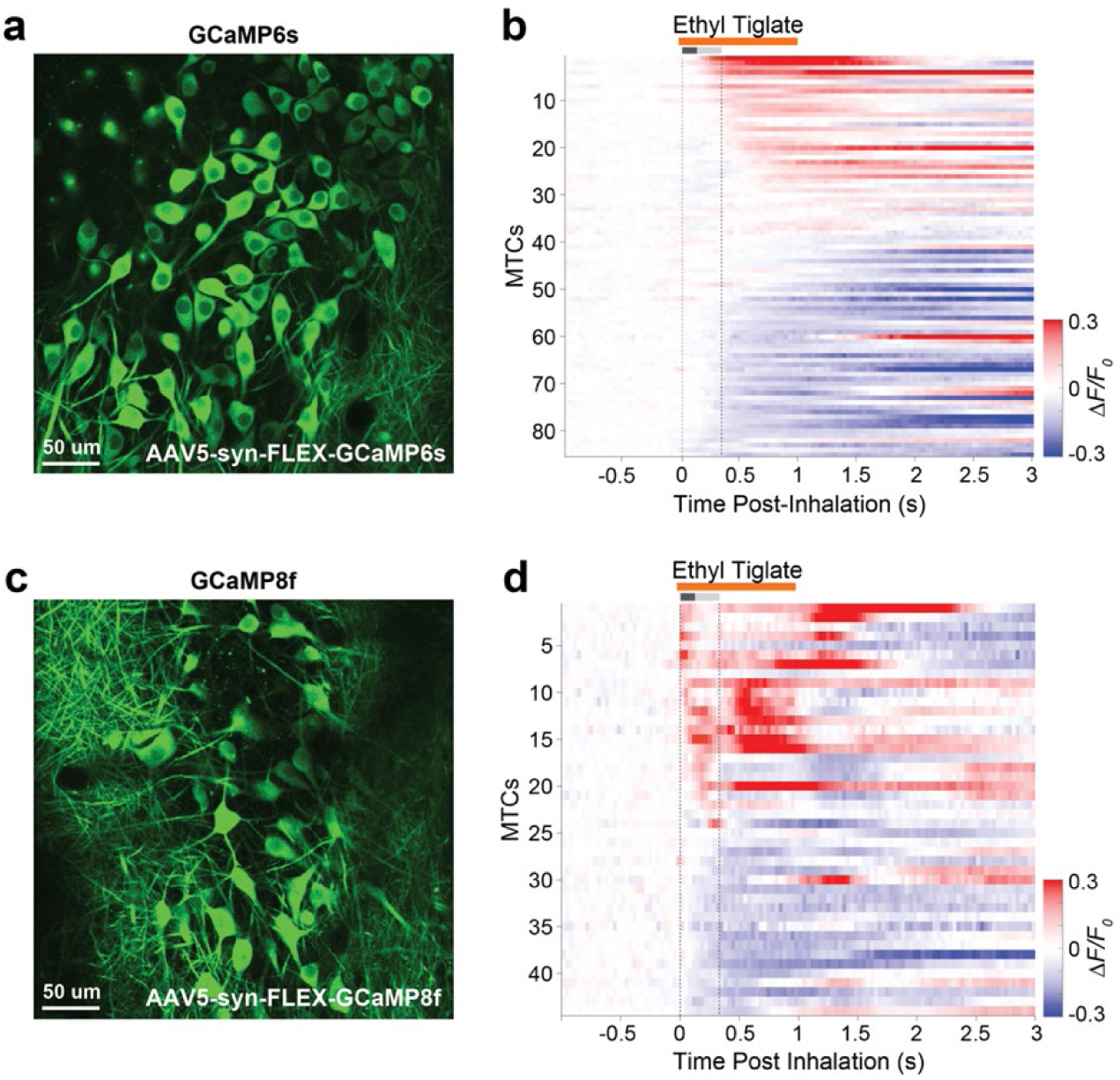
Comparison between GCaMP6s and GCaMP8f performance. **a.** MTCs expressing GCaMP6s (85 MTCs, average of 200 frames). **b.** Mean fluorescence aligned to the onset of inhalation of Ethyl Tiglate (mean of 20 trials, 5% SVP). The first sniff is marked with a gray bar and dotted lines. MTCs were ordered by their response latency. **c.** MTCs expressing GCaMP8f (44 MTCs, average of 200 frames). **d.** Mean fluorescence aligned to the onset of inhalation of Ethyl Tiglate (mean of 15 trials, 5% SVP). The first sniff is marked as in **b**.

**Figure S3.**
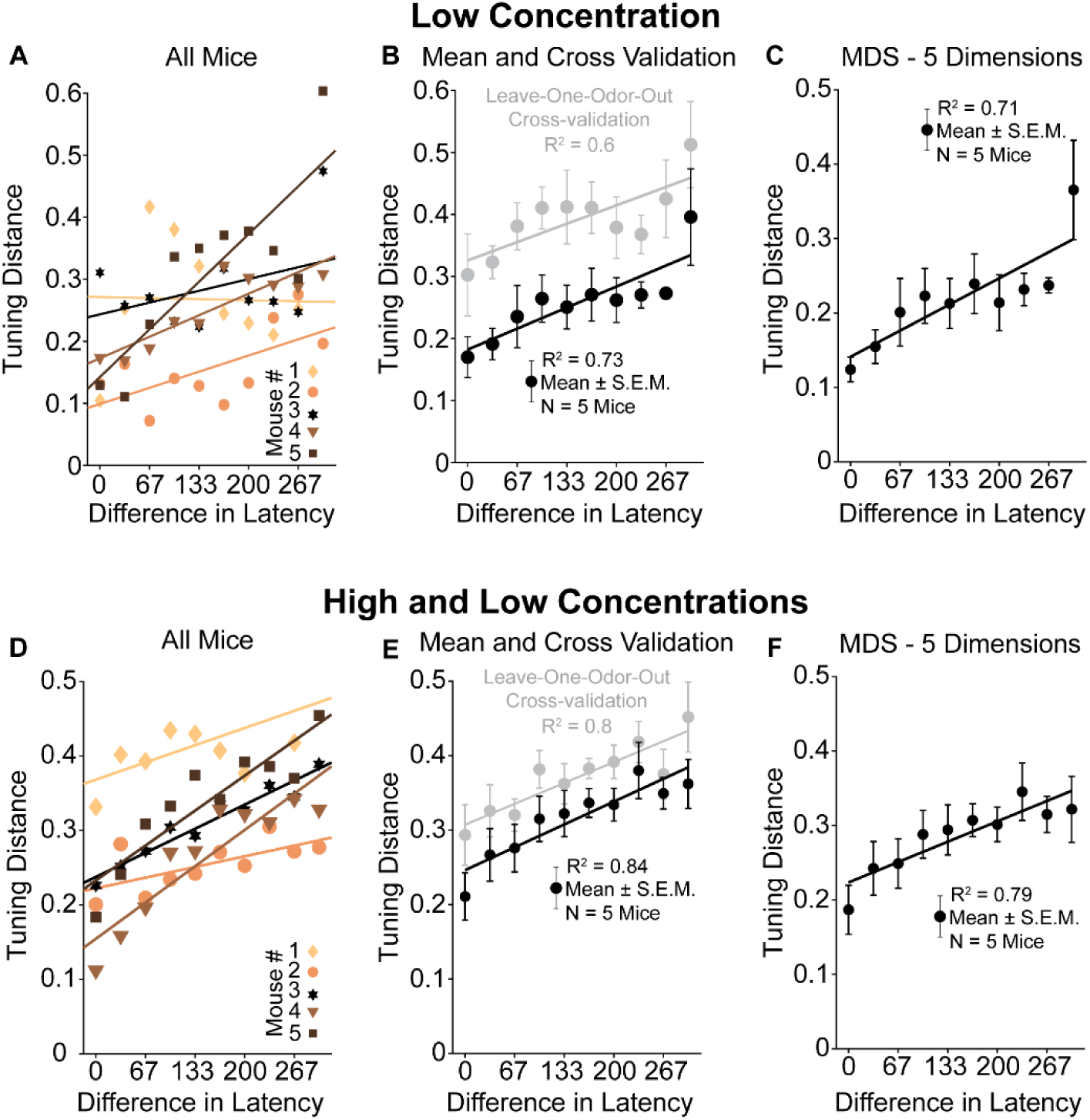
Comparing tuning distance with latency for high and low concentration conditions. **a**. Tuning distance as a function of difference in latency for 5 mice. Only responses to odorants at low concentration used for analyses (8 odorants, 0.5% SVP.) **b**. Mean tuning distance vs. latency across mice (black, slope = 0.017 ms^-1^, R^2^=0.73, p=1.6x10^-3^, 8 odors, 0.5% SVP, n=5 mice, mean +/- SEM) and leave-one-out cross-validation control, where tuning distances were estimated with n-1 odors and the difference in latency was measured for the held-out odor (gray, R^2^ = 0.6, p<0.01, n = 5 mice, mean +/- SEM). **c.** Same as **b** with MTC odor responses projected onto the top 5 MDS dimensions (linear regression, R^2^=0.71, p<0.01, n=5 mice). **d**. Tuning distance as a function of difference in latency for 5 mice. Responses to odorants at both high and low concentrations were used to compute tuning distance and subsequent analyses (8 odorants, 5% and 0.5% SVP., 16 total conditions) **e**. Mean tuning distance vs. latency across mice (black, slope = 0.015 ms^-1^, R^2^=0.84, p=2.1x10^-4^, 8 odors, 5% and 0.5% SVP, n=5 mice, mean +/-SEM) and leave-one-out cross-validation control, where tuning distances were estimated with n-1 odors and the difference in latency was measured for the held-out odor (gray, R^2^ = 0.8, p<0.01, n = 5 mice, mean +/- SEM). **f.** Same as **e** with MTC odor responses projected onto the top 5 MDS dimensions (linear regression, R^2^=0.79, p<0.01, n=5 mice, mean +/- SEM).

**Figure S4.**
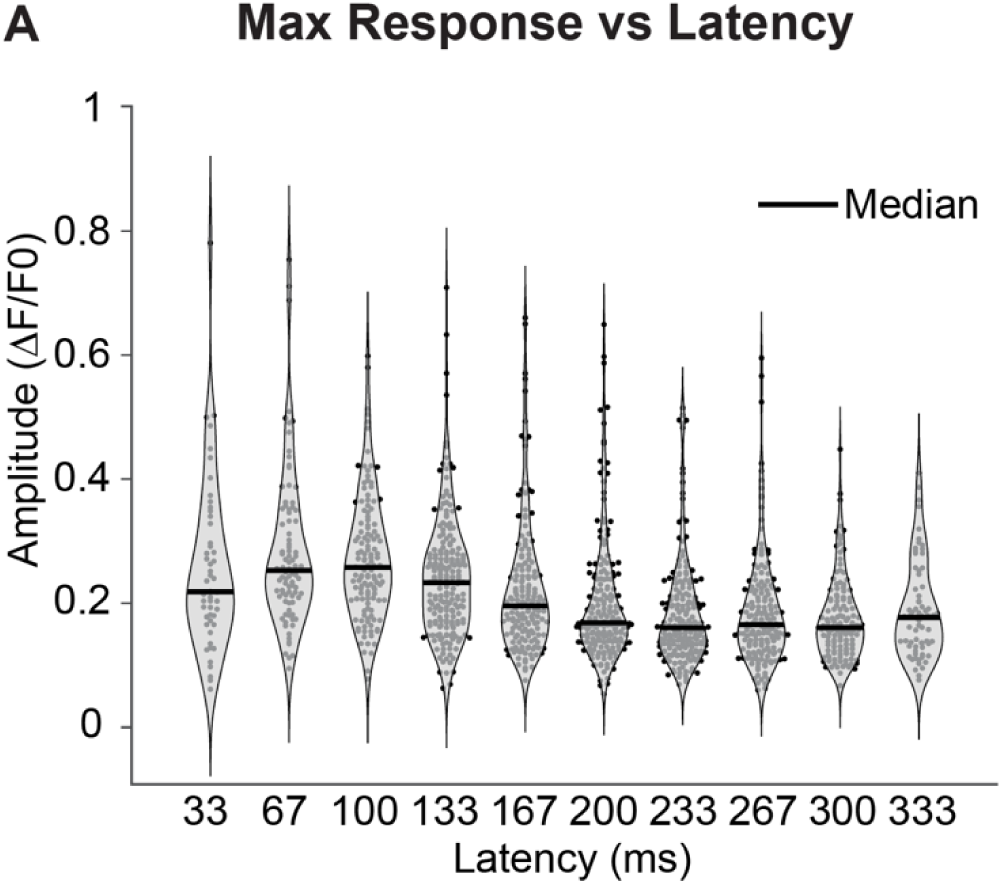
Comparing peak amplitude vs. latency of excitatory odor responses. Violin plot of the peak amplitude of mean excitatory odor responses across 1,376 cell-odor pairs as a function of the latency of their response from the onset of inhalation (627 MTCs, n=5 mice, 8 odors, 5% SVP, mean of 10 trials). Black bars indicate the median of the peak amplitude distribution at each latency.

